# Pharmacological inhibition of the VCP/proteasome axis rescues photoreceptor degeneration in RHO^P23H^ rat retinal explants

**DOI:** 10.1101/2021.06.14.448364

**Authors:** Merve Sen, Oksana Kutsyr, Bowen Cao, Sylvia Bolz, Blanca Arango-Gonzalez, Marius Ueffing

## Abstract

Rhodopsin (*RHO*) misfolding mutations are a common cause of the blinding disease autosomal dominant retinitis pigmentosa (adRP). The most prevalent mutation, *RHO*^P23H^, results in its misfolding and retention in the Endoplasmic Reticulum (ER). Under homeostatic conditions, misfolded proteins are selectively identified, retained at the ER, and cleared via ER-associated degradation (ERAD) and/or autophagy. Overload of these degradation processes for a prolonged period leads to imbalanced proteostasis and may eventually result in cell death. ERAD of misfolded proteins like RHO^P23H^ includes the subsequent steps of protein recognition, targeting for ERAD, retrotranslocation, and proteasomal degradation. In the present study, we investigated and compared pharmacological modulation of ERAD at these four different major steps. We show that inhibition of the VCP/proteasome activity favors cell survival and suppresses P23H-mediated retinal degeneration in RHO^P23H^ rat retinal explants. We suggest targeting this activity as a therapeutic approach for patients with currently untreatable adRP.

## 2. Introduction

Proteostasis requires a complex network of cellular factors, including protein synthesis, folding, and degradation [1]. Protein synthesis is not an error-free process, and approximately 5% of the newly translated proteins contain a sequence error, which can induce protein misfolding and aggregation [2]. Cells are equipped with different physiological mechanisms that prevent or deal with stress caused by misfolded proteins. However, a prolonged imbalance between the folding demand and the cell folding capacity will disrupt cellular fitness, cause disease phenotypes, eventually lead to cell death.

Retinitis Pigmentosa (RP) is a group of inherited retinal degeneration (IRD) characterized by the progressive loss of photoreceptor cells that ultimately leads to blindness [3,4]. In autosomal dominant RP (adRP), most common genetic factors are linked to point mutations in the rhodopsin (*RHO*) gene, with known biochemical defects affecting synthesis, folding, or transport of RHO to the outer segment (OS) and initiate disease despite the presence of wild-type (WT) RHO protein [5-8]. Since RHO is the most abundant protein in photoreceptors—making up 25% of total rod cell protein [6]— it is easy to understand how mutations inducing misfolding of this protein will quickly overload the cellular folding capacity.

The substitution of proline 23 by histidine (*RHO*^*P23H*^) is the most common point mutation associated with adRP in North America [9]. This mutation results in the improper folding of the protein, enhancing its retention in the ER and thereby promoting the formation of intracellular aggregates [10,11]. RHO misfolding and ER retention can activate the unfolded protein response (UPR), aiming to reduce RHO translation, induce expression of chaperones involved in RHO folding, and activate degradation of misfolded RHO [12,13].

Misfolded RHO^P23H^ is retained in the ER, and 90% undergoes extensive degradation via ER-associated protein degradation (ERAD) and/or autophagy [10,14]. As schematized in Figure 1, in ERAD, misfolded RHO^P23H^ is first recognized by molecular chaperones (such as heat shock protein 90 –Hsp90) to subject it to additional folding cycles. If the refolding process fails, misfolded RHO is targeted to the ER Mannosidase 1 (ERM1), which trims terminal mannoses of glycoproteins, preventing them from becoming permanently trapped in the reglucosylation/folding cycle [15]. ERM1 inclines misfolded RHO for the retrotranslocation and targets it for ERAD by cooperating with ER degradation-enhancing α-mannosidase-like 1 (EDEM1). Misfolded RHO is ubiquitinated, and the cytoplasmic valosin-containing protein (VCP) is recruited to the retrotranslocation site within the ER membrane. Ubiquitinated RHO interacts with VCP, is retrotranslocated into the cytosol, and is then delivered to the proteasome for its degradation [15,16] (Figure 1).

**Figure 1:**
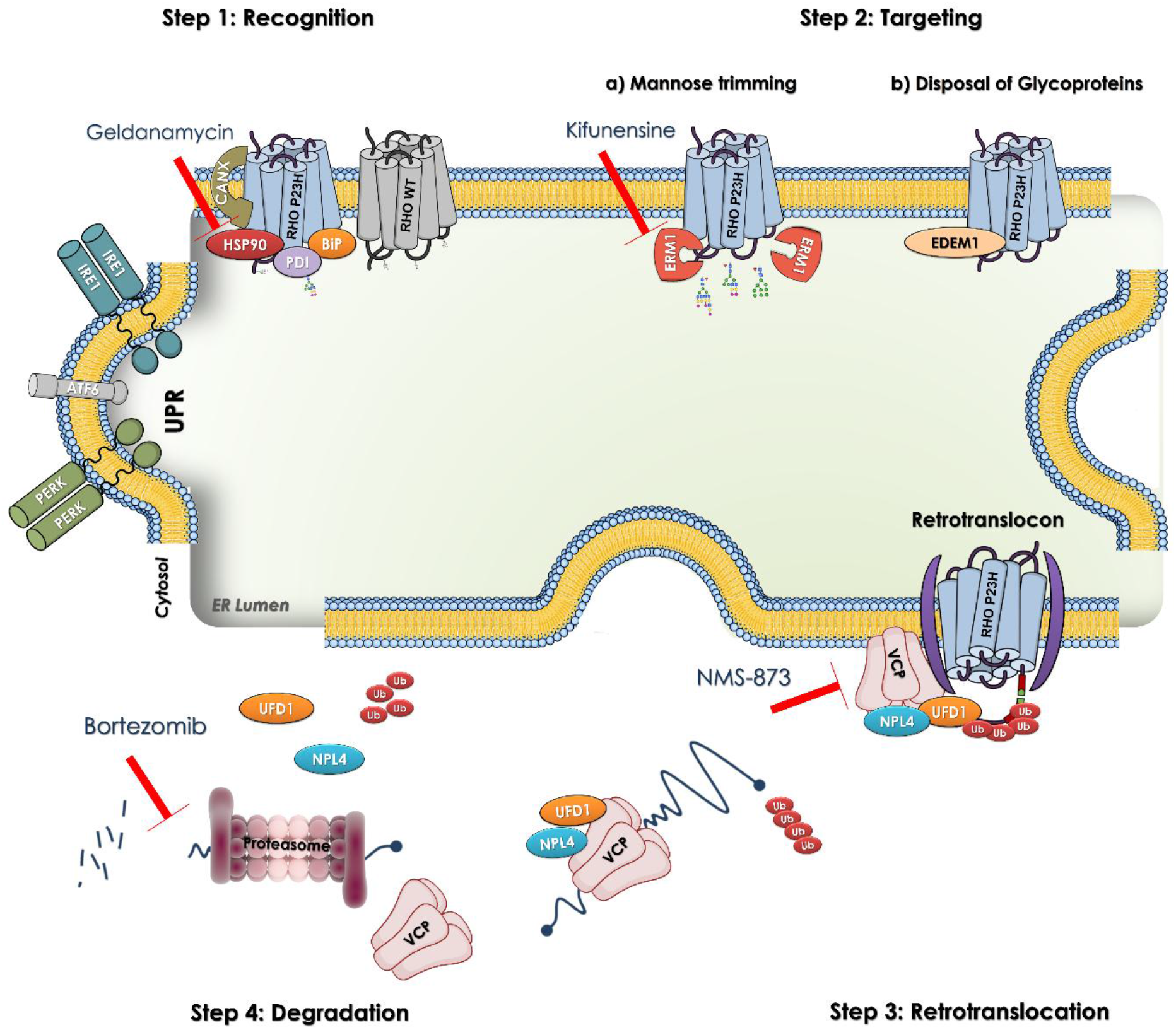
Schematic illustration of ERAD of misfolded RHO^P23H^. **Step 1: RHO recognition**. RHO^P23H^ (blue) is a misfolded glycoprotein that is first recognized by ER-resident chaperones such as Hsp90, BiP, and PDI to start the refolding cycle. The prolonged presence of these mutant forms in the ER activates the unfolded protein response (UPR). **Step 2: RHO targeting**. (a) After the failure to refold, misfolded RHO is trapped in the quality control/folding cycle and becomes a target for ERM1 to remove mannose residues. (b) After mannose removal, misfolded RHO is targeted to the ERAD retrotranslocation site by EDEM1, a facilitator of glycoprotein disposal from the ER. **Step 3: RHO retrotranslocation**. RHO^P23H^ is ubiquitinated and presented to the VCP-Ufd1-Npl4 complex, recruited to the retrotranslocation site by the ER membrane proteins, and retrotranslocated into the cytosol. **Step 4: RHO proteasomal targeting and degradation**. The VCP complex delivers mutant RHO proteins to the proteasome for their degradation. Red arrows show different inhibition targets to modulate distinct ER-related executers. Geldanamycin is an inhibitor for Hsp90, Kifunensine blocks ERM1, NMS-873 is a VCP inhibitor, and Bortezomib interrupts the function of the proteasome. Abbreviations: ERAD: Endoplasmic reticulum-associated degradation, RHO: rhodopsin, WT: wild-type, Hsp90: Heat shock protein 90, PDI: Protein disulfide isomerase, BIP: Binding immunoglobulin protein, ERM1: Endoplasmic reticulum mannosidase 1, EDEM1: ER degradation-enhancing α-mannosidase-like 1, VCP: Valosin - containing protein, Npl4: Nuclear protein localization 4, Ufd1: Ubiquitin fusion degradation, Ub: Ubiquitin, UPR: Unfolded protein response, IRE1: Inositol-requiring protein 1, ATF6: Activating transcription factor 6, PERK: Protein kinase RNA-like ER kinase. This schematic model is inspired and changed from [15,16].

Pharmacological interventions of ERAD effectors can reduce cell death, improve protein folding and enhance RHO trafficking [6,10,11,17]. Inhibition of VCP or proteasome activity strongly slows down photoreceptor degeneration in *Drosophila* induced by *Rh1*^*P37H*^ (the equivalent of mammalian P23H) [14]. We have recently shown that pharmacological inhibition or gene silencing of VCP confer neuroprotection in RHO^P23H^ transgenic rats and knock-in mice [17,18]. Moreover, Hsp90 inhibition enhances visual function, delays photoreceptor degeneration in RHO^P23H^ rats, and reduces the intracellular accumulation in neuroblastoma cells transfected with RHO mutation R135L [19].

In the present study, we investigated and compared the pharmacological modulation effects of ERAD at four different major steps. These pathways were modulated using specific inhibitors in well-characterized RHO^P23H^ transgenic rat organotypic retinal cultures. We utilized Geldanamycin (GA) to inhibit the protein-folding mechanism in the ER by interrupting Hsp90 activity, the mannosidase inhibitor Kifunensine (KIF) to block the processing of glycoproteins by ERM1, NMS-873 to interfere with the retrotranslocation of protein by inhibiting VCP ATPase activity, and the 26S proteasome inhibitor Bortezomib (BO) to suppress proteasomal degradation. We show here that targeting the VCP/proteasome activity favors cell survival and suppresses degeneration in RHO^P23H^ rat retinal explants. We suggest targeting this activity as a therapeutic approach for patients with currently untreatable adRP.

## 3. Results

To analyze the impact of interference with different steps of ERAD on photoreceptor degeneration, we employed primary retinal culture from RHO^P23H^ transgenic rats as a serum-free organotypic culture system preserving the entire mounted flat retina with its adherent RPE and the retinal ganglion cell layer uppermost [20]. RHO^P23H^ transgenic rats, a genetically engineered RHO mutant model, closely mimic the expression pattern of the disease in humans. The RHO^P23H^ rat presents an early and quick progression with a degeneration peak in heterozygous animals at postnatal day (PN) 15 [21]. We explanted RHO^P23H^ rat retinas at PN9 and cultured them for six days until the peak of degeneration at PN15. Photoreceptors undergoing cell death were identified by the terminal deoxynucleotidyl transferase dUTP nick end labeling (TUNEL) assay and calculation of the percentage of positive cells compared to the total outer nuclear layer (ONL) cell nuclei.

### Hsp90 or ERM1 inhibition is detrimental for RHO^P23H^ rat retinae ex vivo

Recognition and targeting of misfolded proteins are the two first steps, determining their fate in the ERAD process. Therefore, we initially investigated whether the inhibition of key elements in these two steps will affect the degenerative course in the RHO^P23H^ retinae. To inhibit the first step of protein recognition, we selected Geldanamycin (GA), which binds competitively with ATP to the N-terminal ATP-binding site of Hsp90 [22]. As an inhibitor for the second step of mannose trimming, we used Kifunensine (KIF), a class I α-mannosidase inhibitor, to inhibit ERM1 [23,24]. Organotypic retinal cultures allowed us to screen different GA and KIF doses, which were chosen according to several *in vitro* assays as reported (Table S1), to inhibit Hsp90 and ERM1 proteins, respectively. Following treatment of the organotypic retinal cultures with either GA or KIF, we evaluated the effect by calculating the percentage of TUNEL positive cells in the ONL of treated RHO^P23H^ retinae.

GA treatment at 0.01 µM and 0.1 µM for 6 days did not protect the retinas. No significant effect in the percentage of TUNEL-positive cells in the ONL of RHO^P23H^ rats was observed. Contrary, retinae treated with 1 µM GA exhibited increased degeneration with a significant increase in the number of dying photoreceptor cells (Vehicle: 3.844 % ± 1.2; 1 µM GA: 8.313 % ± 2.7, p<0.05; Figure 2).

**Figure 2:**
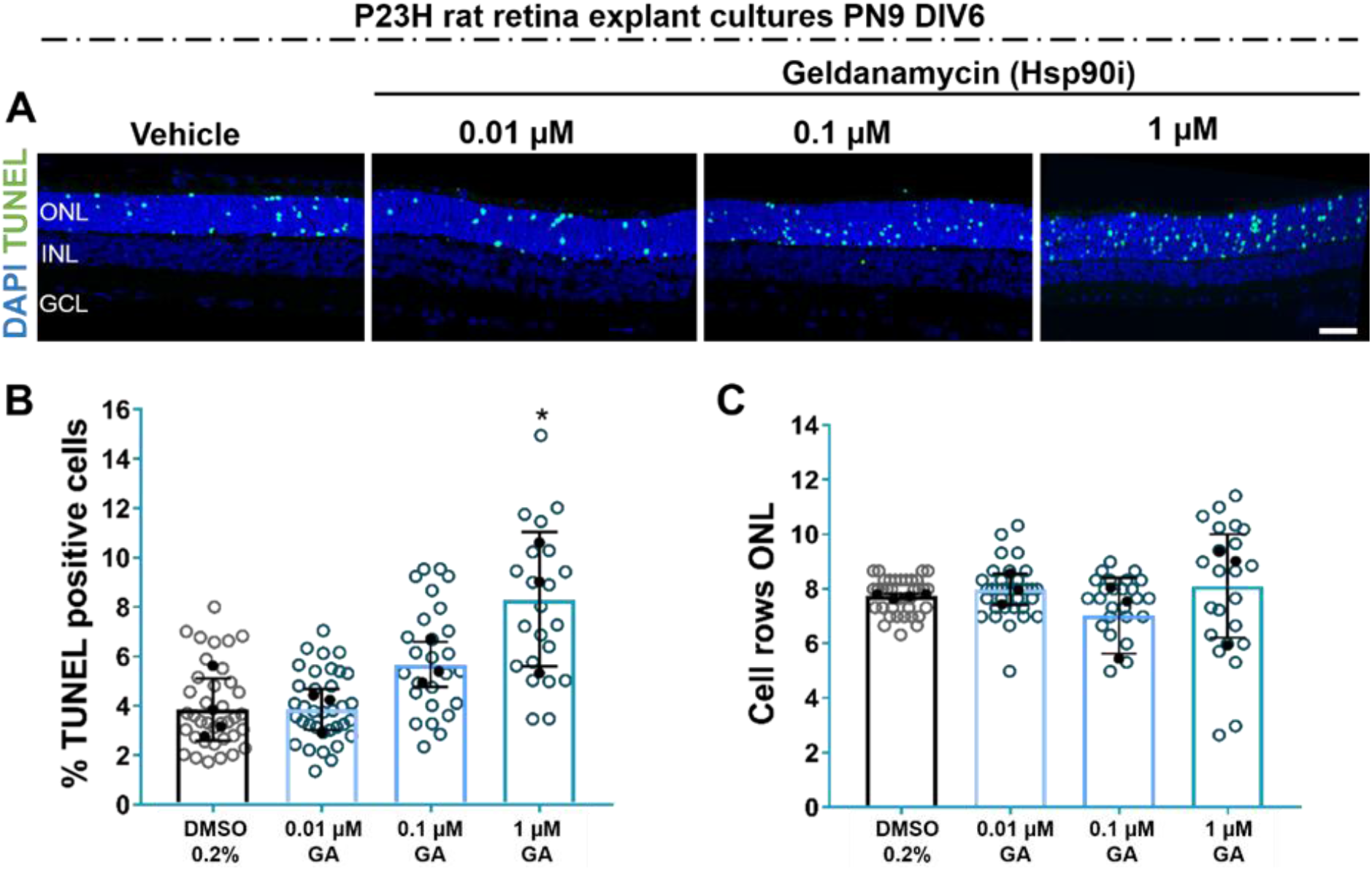
Inhibition of Hsp90 does not improve photoreceptor cell survival in RHO^P23H^ retinae *in vitro*. Retinae from RHO^P23H^ transgenic rats were explanted at postnatal day 9, cultivated for 6 days (PN9 DIV6), and treated every second day with different concentrations Geldanamycin (GA, Hsp90 inhibitor). 0.2% DMSO was used as vehicle control. (A) Explants were stained with TUNEL assay to differentiate photoreceptors undergoing cell death (green) using nuclei counterstaining with DAPI (blue). (B) Bar chart shows the percentage of TUNEL-positive cells in the ONL. A significant increase in the percentage of dying cells was observed after treatment with 1µM GA compared to the respective control. (C) Comparison of the number of remaining cell rows in the ONL. Retinae treated with Hsp90 inhibitor did not show any significant change in the number of photoreceptor cell rows. Scale bar is 50 µm. The values were quantified by scoring several images (circles) from DMSO-treated retinae (closed circles, n=4) and GA-treated retinae at 3 different concentrations (0.01, 0.1, and 1 µM, closed circles, n=3 for each). The data are presented as mean ±SD, and one-way ANOVA analysis was performed. *p<0.05. Hsp90i: Hsp90 inhibitor, ONL: outer nuclear layer, INL: inner nuclear layer, GCL: ganglion cell layer, GA: Geldanamycin.

We next evaluated the role of mannose trimming in P23H photoreceptor degeneration by using the inhibitor KIF. Similarly to GA, retinae treated with 10 or 100 µM KIF showed induced cell death, detected by an increased percentage of TUNEL-positive cells in the ONL (Vehicle: 4.149 % ± 0.8; 10 µM KIF: 5.674 % ± 0.5 p<0.05; and 100 µM KIF: 6.579 % ± 0.2 p<0.001; Figure 3). Treatment with the lower dose of 1 µM KIF did not show any effect.

**Figure 3:**
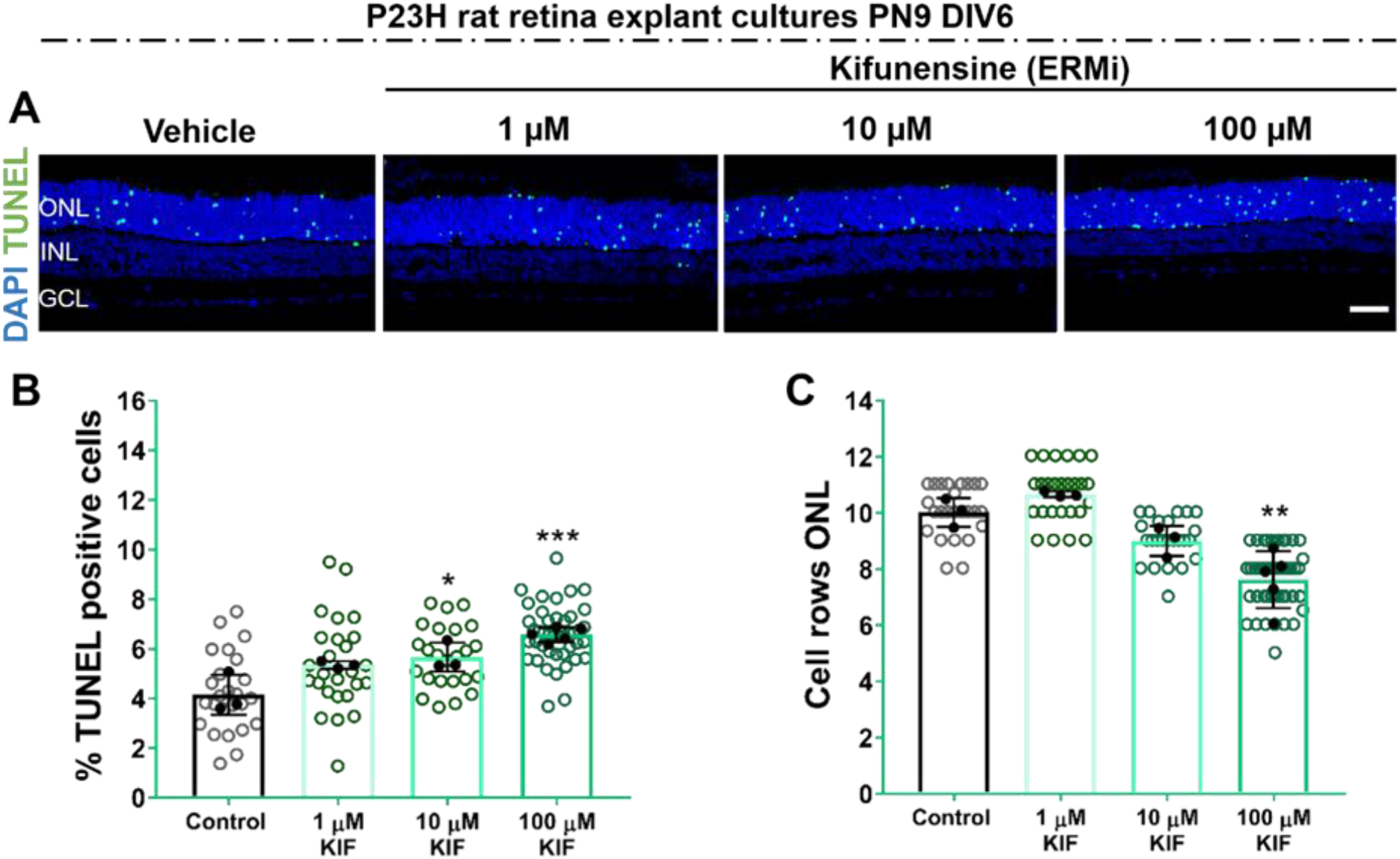
Pharmacological inhibition of Kifunensine (KIF) increases cell toxicity in photoreceptor cells in RHO^P23H^ rats *in vitro*. Retinae from RHO^P23H^ transgenic rats were explanted at postnatal day 9, cultivated for 6 days (PN9 DIV6), and treated every second day with different concentrations of the ERM1 inhibitor KIF or vehicle control. (A) Explants were stained with TUNEL assay to differentiate photoreceptors undergoing cell death (green) using nuclei counterstaining with DAPI (blue). (B) Bar chart shows the percentage of TUNEL-positive cells in the ONL. A significant increase in the percentage of dying cells was observed after the KIF treatment. (C) Comparison of the number of remaining cell rows in the ONL. Retinae treated with KIF showed a significant decrease in the number of photoreceptor cell rows. Scale bar is 50 µm. Values were quantified by scoring several images (open circles) from 3 retinae (closed circles, n=3) per treatment for control, 1 and 10 µM KIF, and 5 retinae for 100 µM KIF (n=5). The data are presented as mean ±SD, and one-way ANOVA analysis was performed. ***p<0.001, **p<0.01, *p<0.05. ERMi: Endoplasmic reticulum mannosidase inhibitor, ONL: outer nuclear layer, INL: inner nuclear layer, GCL: Ganglion cell layer, Kifunensine: KIF.

**Figure 4:**
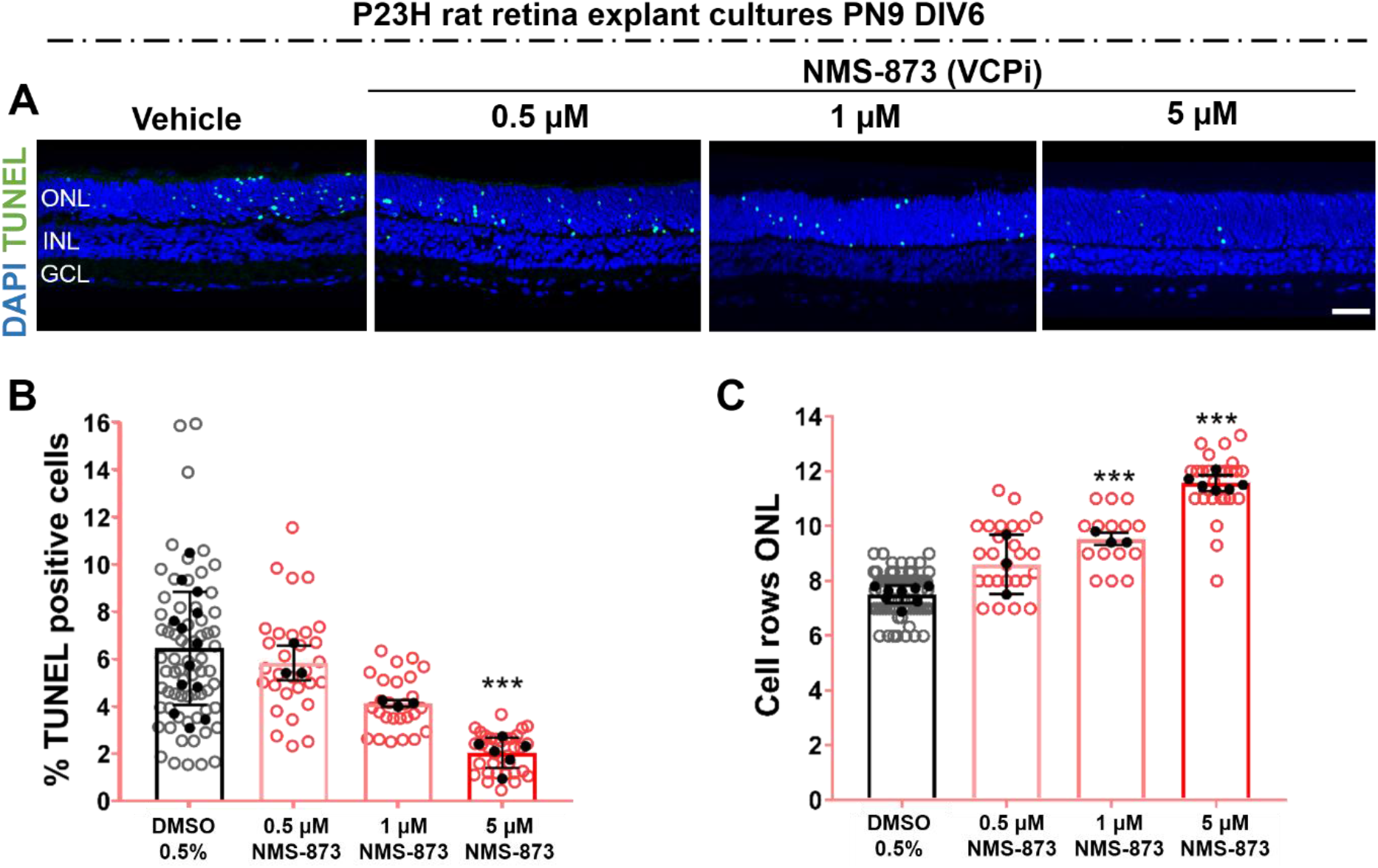
VCP inhibition significantly reduced the number of dying cells and increased cell survival in RHO^P23H^ rats in vitro. Retinae from RHO^P23H^ transgenic rats were explanted at postnatal day 9, cultivated for 6 days (PN9 DIV6), and treated every second day with different concentrations of the VCP inhibitor -NMS-873. 0.5% DMSO was applied as corresponding vehicle control. (A) TUNEL assay reveals photoreceptors undergoing cell death (green), nuclei were counterstained with DAPI (blue). (B) Bar chart shows the percentage of TUNEL-positive cells in the ONL. A decrease in the percentage of dying cells was observed after NMS-873 treatment in a dose-dependent manner, significant at 5µM. (C) Comparison of the number of cell rows in the ONL. Retinae treated with VCP inhibitor NMS-873 showed higher numbers of photoreceptor cell rows. Scale bar is 50 µm. Values were quantified by scoring several images (open circles) from 9 retinae (closed circles, n=13 for TUNEL assay, n=9 for cell rows) per treatment for DMSO 0.5 %, 3 retinae (n=3) for 0.5 and 1 µM NMS-873 and 6 retinae (n=6) for 5 µM NMS-873. The data are presented as mean ±SD, and one-way ANOVA analysis was performed at *p<0.05, **p<0.01, ***p<0.001, ****p<0.0001. VCPi: VCP inhibitor, ONL: outer nuclear layer, INL: inner nuclear layer, GCL: Ganglion cell layer.

As a second measurement to test for potential protective effects of HSP90 or ERM1 inhibition on photoreceptor survival, we next counted photoreceptor cell rows in the ONL. We found that GA did not affect the number of remaining photoreceptor cell rows at the evaluated concentrations. However, the observed increase in TUNEL-positive cells with 100 µM KIF was reflected by a significantly reduced number of remaining cell rows in the ONL of KIF-treated RHO^P23H^ retinae (Vehicle: 10.03 rows ± 0.5; 100 µM KIF: 7.628 rows ± 1.0 p<0.01; Figure 3).

These results indicate that inhibition of pathways involved in recognizing misfolded RHO or targeting RHO^P23H^ to ERAD by the inhibitors GA or KIF is detrimental and accelerates photoreceptor cell loss in RHO^P23H^ rat retinal explants.

### VCP inhibition enhances photoreceptor cell survival in RHO^P23H^ retinal cultures

Following recognition and targeting, RHO^P23H^ ERAD substrates are retrotranslocated across the ER membrane into the cytosol. In fact, for almost all ERAD substrates, VCP has an essential role in substrate retrotranslocation in yeast and mammals [25]. VCP extracts substrates from the ER to the cytosol in an ATP-dependent manner (retrotranslocation). Ubiquitin-labeled such substrates are then directed to the proteasome [26]. Previous studies in cellular and *Drosophila* models indicate that excessive retrotranslocation activity causes cell loss in the presence of mutant *RHO*^*P23H*^ or *Rh1*^*P37H*^, respectively [10,14]. Also, VCP inhibition has been shown to be protective in RHO^P23H^ animal models [17].

Here, we compare inhibition of VCP-dependent retrotranslocation with the interference at different other ERAD steps under the same experimental conditions in RHO^P23H^ rat retinal cultures. To test the impact of VCP inhibition, we employed the allosteric non-ATP-competitive VCP inhibitor NMS-873, one of the most potent and specific VCP inhibitors described to date, at three different doses. NMS-873 alters the binding affinity of the D1–D2 interdomain linker of VCP and inhibits ATPase activity [27]. When applied to RHO^P23H^ rat retinal cultures under the same experimental conditions as described above, NMS-873 reduced photoreceptor cell death in a dose-dependent manner, showing minimum cell death and maximum photoreceptor cell protection at 5 µM. The percentage of TUNEL-positive cells decreased (Vehicle: 6.760 % ± 2.6; 5 µM NMS-873: 2.043 %± 0.6 p<0.001, Figure 5A-B) and the number of photoreceptor cell rows significantly increased (Vehicle: 7.25 rows ± 0.3; 0.5 µM NMS-873: 8.604 rows ± 1.0 p<0.05, 1 µM NMS-873: 9.533 rows ± 0.2 p<0.01; and 5 µM NMS-873: 11.56 rows ± 0.2 p<0.001; Figure 5 A-C).

**Figure 5:**
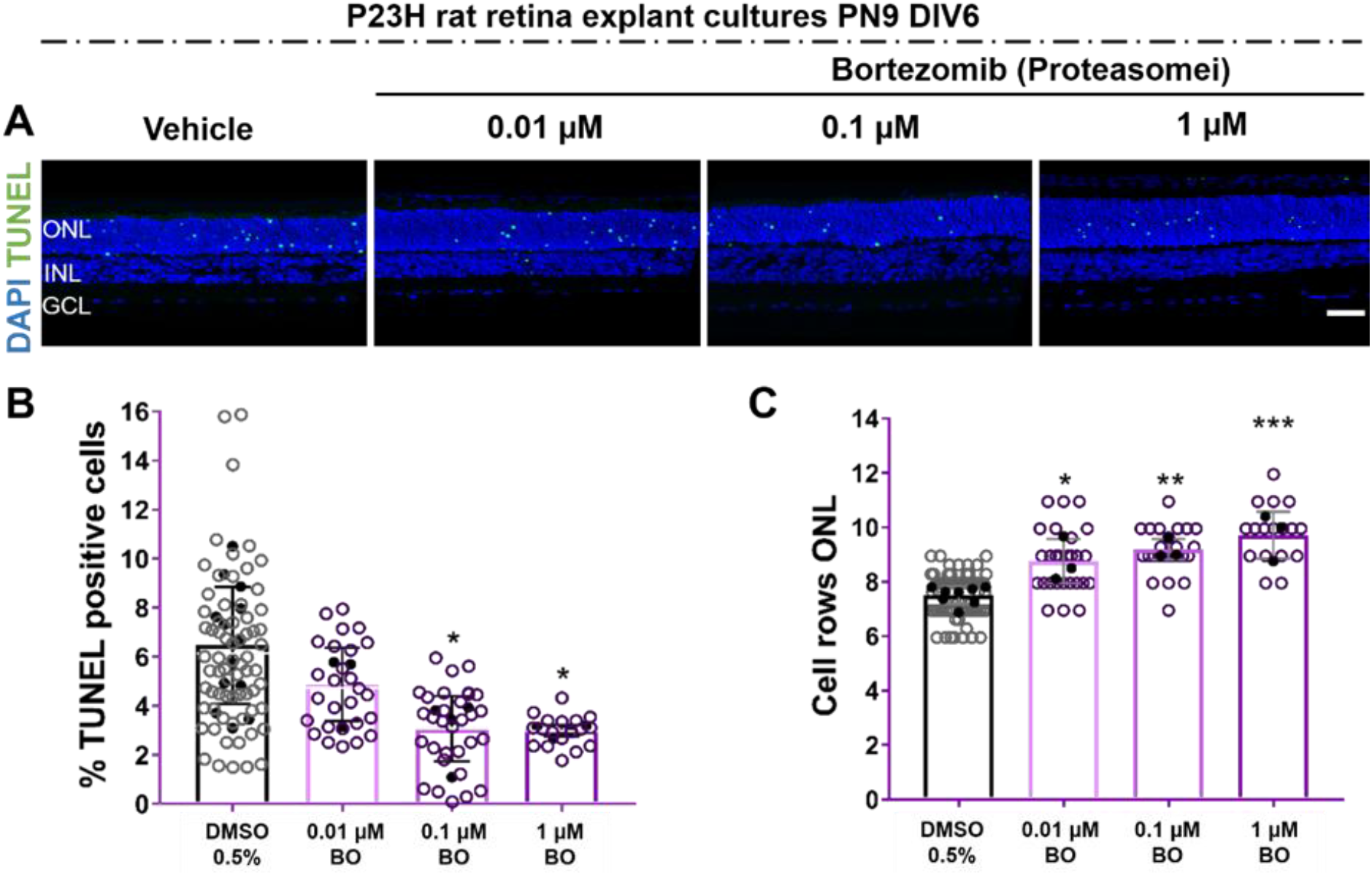
Proteasome inhibition by Bortezomib (BO) enhances photoreceptor cell survival in RHO^P23H^ rat retinae *in vitro*. Retinae from RHO^P23H^ transgenic rats were explanted at postnatal day 9, cultivated for 6 days (PN9 DIV6), and treated every second day with different concentrations of proteasome inhibitor -BO. 0.5% DMSO was used as corresponding vehicle control. (A) Explants were stained with TUNEL assay to differentiate photoreceptors undergoing cell death (green) using nuclei counterstaining with DAPI (blue). (B) Bar chart shows the percentage of TUNEL-positive cells in the ONL. A significant decrease in the percentage of dying cells was observed after BO treatment. (C) Comparison of the number of remaining cell rows in the ONL. Retinae treated with BO showed an increase in the number of photoreceptor cell rows. Scale bar is 50 µm. Values were quantified by scoring several images (open circles) from 3 retinae (closed circles, n=3) per treatment for DMSO and 0.01 µM BO, 4 retinae (n=4) for 0.1 and 1 µM BO. The data are presented as mean ±SD, and one-way ANOVA analysis was performed at **p<0.01, *p<0.05. Proteasomei: Proteasome inhibitor, ONL: outer nuclear layer, INL: inner nuclear layer, GCL: Ganglion cell layer, BO: Bortezomib.

### 26S Proteasome inhibition reduces photoreceptor degeneration in RHO^P23H^ rat explants

VCP activity and proteasomal activity are two interacting elements of substrate degradation during ERAD [10]. Proteasome overload has been previously shown to contribute to photoreceptor cell death in RHO^P23H^ retinae [14,28].

To investigate if proteasome inhibition could confer a protective effect in RHO^P23H^ rat retinal explants, we used the inhibitor Bortezomib (BO). BO blocks proteasome-targeted proteolysis by inhibiting the 26S proteasome, a large protease complex that degrades ubiquitinated proteins [29].

RHO^P23H^ retinal explants were treated with different concentrations of BO compared to DMSO control. As shown in Figure 5, treatment with BO significantly reduced the percentage of cell death (Vehicle: 5.473 % ± 1.04; 0.1 µM BO: 3.066 % ±1.3 p<0.05; and 1 µM BO: 2.993 % ± 0.2 p<0.05). Accordingly, the number of surviving photoreceptor cell rows in the ONL increased (Vehicle: 7.665 rows ± 0.1; 0.1 µM BO: 9.194 rows ± 0.3 p<0.05; and 1 µM BO 9.717 rows ± 0.8 p<0.01), indicating that this treatment conferred neuroprotection (Figure 5 A-C).

Thus, pharmacological inhibition of the VCP/proteasome axis is protective for RHO^P23H^ photoreceptors, further suggesting that excessive retrotranslocation and/or excessive degradation of RHO is a critical detrimental event in the RHO^P23H^ retina.

### Only VCP inhibition enhances RHO trafficking to the outer segment in RHO^P23H^ rat retinal explants

*RHO*^*P23H*^ mutation is characterized by RHO mislocalization even before photoreceptor degeneration starts. Impaired RHO transport is associated with disorganized and shortened photoreceptor OSs [30]. WT control retinae exhibit high RHO expression, and the protein localizes mainly to the photoreceptor’s OS. In contrast, RHO^P23H^ retinae show RHO staining distributed throughout the ONL [17]. Therefore, we investigated whether different ER modulators (GA, KIF, NMS-873, and BO) could reconstitute proper physiological RHO trafficking to the OS.

In RHO^P23H^ corresponding vehicle control retinae, immunofluorescence analysis showed that RHO was mislocalized, showing an accumulation in the ONL (Figure 6). Using this as a reference, we found similar staining after treatment at all evaluated doses of GA, KIF, and BO. Only VCP inhibition by NMS-873 restored the distribution of RHO to the OS in a dose-dependent manner (Figure 6).

**Figure 6:**
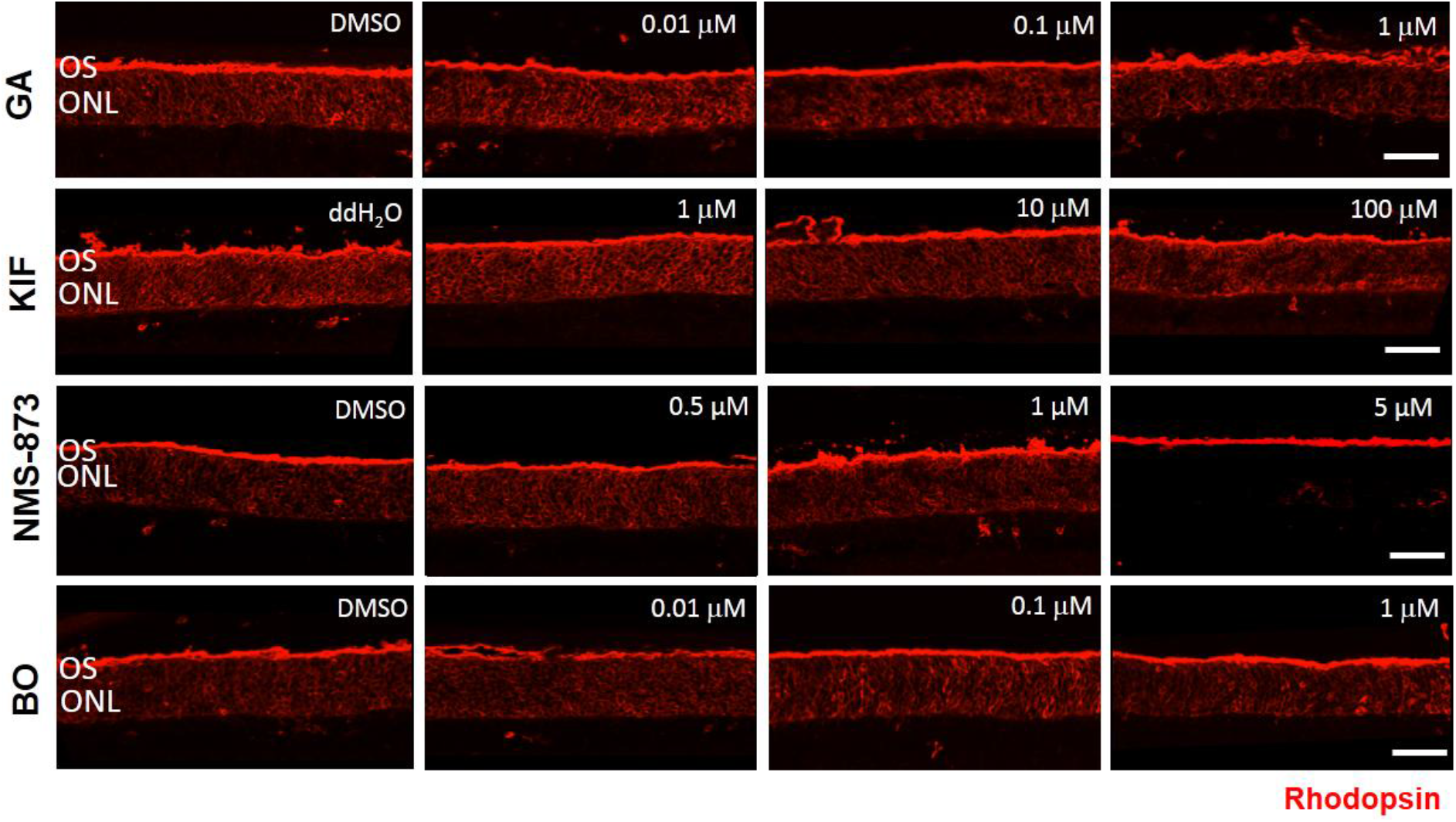
Only VCP inhibition mainly restores Rhodopsin (RHO) localization to the outer segment (OS) in RHO^P23H^ transgenic rat retinae *in vitro*. Fluorescent labeling in cryosections designates the location of RHO (red staining) in RHO^P23H^ retinae organotypic cultures at PN15, explanted at postnatal day 9 and cultivated for 6 days (PN9 DIV6), treated with corresponding vehicle (ddH^2^O or DMSO) or different concentrations of Hsp90 inhibitor GA, ERM1 inhibitor KIF, VCP inhibitor NMS-873 and proteasome inhibitor BO. In the vehicle-treated groups, RHO distribution was observed with intense fluorescence staining in the ONL. Only in retinae treated with 5 µM NMS-873, a reduction of the fluorescence in the ONL is observed. Other inhibitors do not affect RHO distribution. Scale bar is 50 µm. PN: postnatal, DIV: days *in vitro*, OS: outer segment, ONL: outer nuclear layer, GA: Geldanamycin, KIF: Kifunensine, BO: Bortezomib, i: inhibitor.

In conclusion, modulation of ERAD either by inhibition of VCP or 26S proteasome activity decreases photoreceptor cell death and improves photoreceptor survival. However, only VCP inhibition enhances RHO localization to the OS.

## 4. Discussion

Misfolded RHO^P23H^ is retained in the ER and then identified by ER folding and chaperone systems, processed by ERAD, and retrotranslocated to the cytosol for degradation by the proteasome [6]. In this scenario, excessive ERAD combined with aggregation-associated proteotoxicity may exceed the capacity of cellular protein clearance, resulting in acute cellular stress and subsequent processes initiating cellular degeneration [31]. Several studies of pharmacological modulation of the cellular proteostasis network have been performed, looking to rebalance proteostasis and potentially unmask novel therapeutic targets. Here, we evaluated four substances able to interfere at the different ERAD steps and tested them under the same experimental conditions to compare their effects. Thus, we selected one of the most commonly used autosomal dominant RP models, the RHO^P23H^ transgenic rat, using a serum-free retinal organotypic system as a unifying approach, and evaluated cell death, cell survival, and RHO immunostaining distribution, choosing VCP inhibition as a positive reference.

VCP is a key interactor of misfolded RHO^P23H^ that allows its retrotranslocation and proteasomal clearance [10]. We have found that inhibition of VCP or proteasome individually attenuates cell death and preserves photoreceptor structure and function in an insect model of adRP [14] and shown that VCP inhibition [17] or silencing [18] mitigates disease progression in RHO^P23H^ rodent models. Here, we were able to substantiate these previous findings when we tested VCP inhibition by NMS-873 in the RHO^P23H^ transgenic model and confirmed that inhibition of retrotranslocation had a protective effect on degenerating photoreceptors. In addition, we found that the inhibition of 26S proteasome activity also enhanced photoreceptor survival in RHO^P23H^ rat retinal explants, however, without the improved protein trafficking observed with VCP inhibition.

In contrast, inhibition of initial steps in ERAD (recognition and targeting of misfolded RHO) does not increase photoreceptor cell survival. Indeed, our results indicate that this type of modulation may have the opposite effect and accelerate the degeneration process, suggesting that prolongation of protein folding attempts might be less harmful to the cell than excessive degradation. Regulation of the structure and folding of proteins is particularly important for maintaining cellular homeostasis [32]. In the past, several studies have attempted to directly modulate the folding capacity in the cell as a neuroprotective strategy. One of the targets has been the chaperone Hsp90. Hsp90 participates in protein folding and protein degradation, playing a role in the assembly and maintenance of the 26S proteasome [33].

Nevertheless, inhibition of Hsp90 has shown contradictory results. Single oral gavage of the Hsp90 inhibitor HSP990 has improved visual function and delayed photoreceptor degeneration in RHO^P23H^ rats. However, its longer oral gavage adversely affected visual function [19]. Other reports have also implicated that inhibition of Hsp90 induces a deleterious effect in mice [34] and human RPE cells *in vitro* [35] and exhibit retinal toxicity in dogs *in vivo* [36]. Adverse effects have been reported in clinical trials, including ocular toxicity associated with visual disturbances [37]. In line with that, upregulation of Hsp70 and Hsp90 by treatment with Arimoclomol resulted in photoreceptor protection and improved visual function [38], suggesting that potentiation of the heat shock response, not its inhibition, might relate to neuroprotection. This is supported by our results on Hsp90 inhibition with GA, in which photoreceptor degeneration was even increased in a dose-dependent manner.

ERM1 appears to have a major function in enhancing the degradation of misfolded RHO^P23H^ by directing them to ERAD. Inhibition of ERM1 by KIF showed partial inhibition of enhanced degradation of RHO^P23H^ in a neuroblastoma cell line [39]. However, the mannosidase inhibitor failed to increase cell survival in RHO^P23H^ rat retinal explants in our study.

We found that targeting the last steps in ERAD - i.e., the VCP/proteasome axis - can have protective effects on P23H rat retinae. RHO^WT^ is recruited into aggregates by RHO^P23H,^ and these aggregates form a complex with VCP, leading to degradation of RHO^WT^ [10]. Retention of these aggregates in the inner segment of photoreceptors restrains the amount of RHO reaching the OS. In addition, retrotranslocation of misfolded RHO^P23H^ via VCP and their subsequent degradation requires high energy levels (e.g., ATP hydrolysis) [40], and this could lead to photoreceptor cell death upon P23H-mediated adRP. This is why, inhibition of the VCP/proteasome axis might prevent a severe energy imbalance and enhance photoreceptor cell survival [16,41]. However, the overall protection provided by proteasome inhibition is weaker compared to VCP inhibition. In addition, we observed proper localization of RHO only after VCP inhibition but not after proteasome inhibition. One possible explanation is that proteasome inhibition, even though it could compensate for the energy requirement, is unable to decrease the RHO-VCP complex levels, thereby impeding the trafficking of RHO to the OS.

Inhibition of excessive ERAD and proteasome activities could, in addition, promote refolding. Nevertheless, approaches to promote RHO refolding alone, e.g., treatment with AMPK activator metformin [42], retinoids [43,44], overexpression of Calnexin [45] or BiP [46] have proven challenging and indicate that promoting RHO^P23H^ refolding alone does not result in a protective outcome. Metformin-rescued RHO^P23H^ showed better protein trafficking but was still intrinsically unstable, increasing the rod OS’s structural instability with consequently reduced photoreceptor function and increased photoreceptor cell death [42]. Enhanced folding of RHO^P23H^ by overexpression of Calnexin enhanced the proper folding of RHO, but its loss of function showed a lack of effect *in vitro* [45]. Also, the neuroprotection obtained by BIP gene delivery in RHO^P23H^ line-3 rats is more likely due to suppression of photoreceptor cell death rather than to enhanced RHO folding [46].

In conclusion, we here report that that pharmacological modulation of the VCP/proteasome axis enhances photoreceptor cell survival and preserves retinal structure in the RHO^P23H^ rat model of adRP *in vitro*, presenting a potential therapeutic strategy for adRP. In contrast, other proteostasis regulators in ERAD that control protein integrity or folding did not show any protective effects. Our study highlights the importance of further investigations to accurately characterize the mechanisms regulating protein folding, quality control, and degradation of RHO for the development of new therapeutic approaches. Furthermore, the importance of understanding these processes may not be limited to P23H-mediated retinal degeneration, but may be extendable to several other diseases caused by inefficient protein folding.

## 5. Materials and Methods

### Study approval and animals

Procedures were approved by the Tuebingen University committee on animal protection (§4 registrations from 01.02.2017 and 26.10.2018 AK 15/18 M) and performed in compliance with the Association for Research in Vision and Ophthalmology ARVO Statement on animal use in ophthalmic and vision research. All efforts were made to minimize the number of animals used and their suffering. Homozygous RHO^P23H^ transgenic rats (produced by Chrysalis DNX Transgenic Sciences, Princeton, NJ) of the line SD-Tg(P23H)1Lav (P23H-1) were kindly provided by Dr. M. M. LaVail (University of California, San Francisco, CA) or by the Rat Resource and Research Center (RRRC) at the University of Missouri. To mimic the genetic background of adRP, we used heterozygous P23H rats obtained by crossing homozygous RHO^P23H^ rats with RHO^WT^ rats (CDH IGS Rat; Charles River, Germany). Animals were housed in the Institute for Ophthalmic Research animal facility under standard white cyclic lighting, with access to food and water.

### Organotypic Retinal Explant Cultures of P23H heterozygous transgenic rats

Retinae were isolated with the retinal pigment epithelium (RPE) attached as described previously [20]. Briefly, PN9 and PN20 animals were sacrificed, the eyes were enucleated in an aseptic environment and pretreated with 12% proteinase K (MP Biomedicals, 0219350490) for 15 minutes at 37 °C in R16 serum-free culture medium (Invitrogen Life Technologies, 07490743A). The enzymatic digestion was stopped by 20 % fetal bovine serum (Sigma-Aldrich, F7524). Retina and RPE were dissected, and four radial cuts were made to flatten it. The tissue was transferred to a 0.4 µm polycarbonate membrane (Corning Life Sciences, CLS3412) with the RPE facing the membrane. The retinal explants were cultured in 1 mL of serum-free culture medium consisting of Neurobasal A (NA, Gibco, Carlsbad, CA, USA) supplemented with 2% B-27 supplement (Gibco, Carlsbad, CA, USA), 1% N2 supplement (Invitrogen, Carlsbad, CA, USA), 1% penicillin solution (Gibco, Carlsbad, CA, USA), and 0.4% GlutaMax (Gibco, Carlsbad, CA, USA). Explants were maintained at 37°C in a humidified 5% CO2 atmosphere. The eyes were then randomly assigned to either vehicle control or inhibitors. For each inhibitor, vehicles (either water or DMSO) were applied according to the highest concentration of the inhibitors. Geldanamycin (Calbiochem, 345805) was dissolved in DMSO, and different concentrations of Geldanamycin were applied in 1 mL culture medium (0.01 µM, 0.1 µM, and 1 µM). 0.2% DMSO was used as vehicle control. Kifunensine (Calbiochem, 422500) was dissolved in hot water (55 °C), and different concentrations of drugs were applied in 1mL culture medium (1 µM, 10 µM, and 100 µM). ddH_2_O was applied as corresponding vehicle control. NMS-873 (Xcessbio, M60165-b) was dissolved in DMSO, and different concentrations were applied in 1 mL culture medium (0.5 µM, 1 µM, and 5 µM). Bortezomib (Calbiochem, 5.04314.0001) was dissolved in DMSO, and 0.01 µM, 0.1 µM, and 1 µM concentrations of Bortezomib were applied in culture medium. For NMS-873 and Bortezomib, 0.5% DMSO served as vehicle control. The medium was changed every second day. The PN9 cultures were fixed at DIV6, which corresponds to PN15, the peak of degeneration *in vivo* in age-matched RHO^P23H^ heterozygous rats.

### Histology

Tissues were immersed in 4% paraformaldehyde in 0.1 M phosphate buffer (PB; pH 7.4) for 45 minutes at 4°C, followed by cryoprotection in graded sucrose solutions (10%, 20%, 30%) and embedded in cryomatrix (Tissue-Tek® O.C.T. Compound, Sakura® Finetek, VWR, 4583). Radial sections (14 µm thick) were collected, air-dried, and stored at -20 °C.

### TUNEL assay

TUNEL assay [47] was performed using an *in situ* cell death detection kit conjugated with fluorescein isothiocyanate (Roche, 11684795910). DAPI (Vectashield Antifade Mounting Medium with DAPI; Vector Laboratories, H-1200) was used as a nuclear counterstain.

### Immunofluorescence staining

Sections were incubated overnight at 4 °C with rhodopsin mAb (Sigma-Aldrich, MAB5316). Fluorescence immunocytochemistry was performed using Alexa Fluor® 568 conjugated secondary antibody (Invitrogen, A-11031). Negative controls were carried out by omitting the primary antibody.

### Microscopy and cell counting

All samples were analyzed using Zeiss Axio Imager Z1 ApoTome microscope, AxioCam MRm camera, and Zeiss Zen 2.3 software in Z-stack at 20X magnification. For quantitative analysis, positive cells in the ONL of at least three sections per group were manually counted. The percentage of positive cells was calculated, dividing the number of positive cells by the total number of cells in the ONL. Photoreceptor cell rows were assessed by counting the individual nuclei rows in one ONL and averaging the counts. Graphs were prepared in GraphPad Prism 7.05 for Windows.

### Statistics

The evaluation of TUNEL analysis and photoreceptor cell row counts was performed using GraphPad Prism 7.05 for Windows and one-way ANOVA testing, followed by Bonferroni multiple comparisons test.

## Acknowledgments

This study was supported by funds (to M.Ue. and B.A-G) from FFB (Grant PPA-0717-0719-RAD), the Kerstan Foundation, European Union’s Horizon 2020 research and innovation programme under the Marie Skłodowska-Curie (Grant agreement No. 722717 – project OCUTHER), the Maloch Stiftung and the ProRetina Foundation. The animal husbandry personnel at the Universitätsklinikums Tübingen and Norman Rieger are acknowledged for animal care. Dr. Mohamed Ali Jarboui is acknowledged for the preparation of Figure 1. Dr. Ellen Kilger is gratefully acknowledged for language editing and proofreading.

## Supplementary file

**Table S1:**
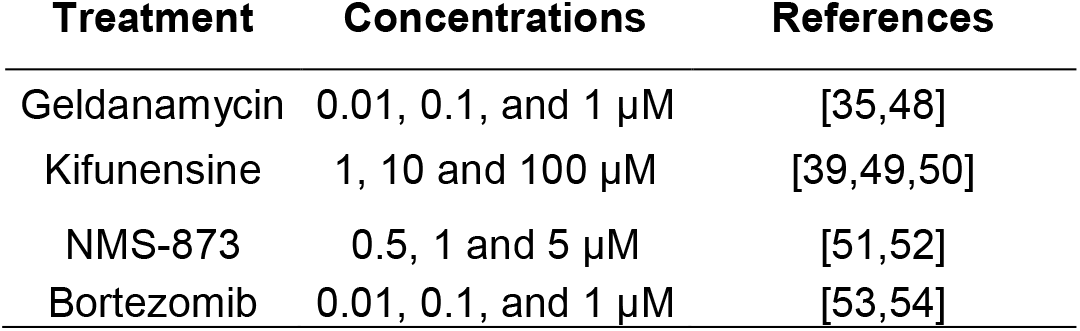
References list for the chosen concentrations in this study.

## Notes

### Competing Interest Statement

The authors have declared no competing interest.

